# Altered Salience Network Connectivity in 6-Week-Old Infants at Risk for Autism

**DOI:** 10.1101/2021.10.27.466195

**Authors:** Tawny Tsang, Shulamite Green, Janelle Liu, Katherine Lawrence, Shafali Jeste, Susan Y. Bookheimer, Mirella Dapretto

## Abstract

Converging evidence implicates disrupted brain connectivity in autism spectrum disorder (ASD); however, the mechanisms linking altered connectivity early in development to the emergence of ASD symptomatology remain poorly understood. Here we examined whether atypicalities in the Salience Network (SN) – an early-emerging neural network involved in orienting attention to the most salient aspects of one’s internal and external environment – may predict the development of ASD markers such as reduced social attention and atypical sensory processing. Six-week-old infants at high-risk for ASD exhibited stronger SN connectivity with sensorimotor regions; low-risk infants demonstrated stronger SN connectivity with prefrontal regions involved in social attention. Infants with higher connectivity with sensorimotor regions had lower connectivity with prefrontal regions, suggesting a direct tradeoff between attention to basic sensory versus socially-relevant information. Early alterations in SN connectivity predicted subsequent ASD symptomatology, providing a plausible mechanistic account for the unfolding of atypical developmental trajectories associated with ASD risk.

---

Shortly after birth, newborns display systematic preferences for faces,^1^ voices,^2^ and biological motion.^3^ These early social-orienting mechanisms are foundational for normative social development.^4^ However, the salience of socially-relevant information appears disrupted in autism spectrum disorders (ASD). Infants who develop ASD show altered developmental trajectories^5^ characterized by reduced attention to social information^6,7^ and heightened awareness of non-social sensory input.^8^ Atypicalities in social versus nonsocial attention are broadly recognized as a marker of ASD risk^9^ and likely contribute to the emergence of social impairments and the restrictive and repetitive behaviors characteristic of ASD.^7,10^ The neurobiological mechanisms underlying these early attentional abnormalities that give rise to autism-related symptoms remains unknown.

Examining early brain connectivity offers a promising lens for investigation.^11^ Indeed, most ASD risk genes impact synapse formation and function, presenting a biological pathway for ASD that converges on brain connectivity. Neuroimaging studies have consistently implicated atypical brain network dynamics in ASD,^11^ with recent evidence demonstrating that 6-month-olds who later develop ASD already exhibit systematic differences in whole-brain functional connectivity.^12^ Thus, early deviations in brain network connectivity may provide a biomarker of ASD risk prior to the emergence of behavioral symptoms.

One early emerging functional brain network of particular interest in understanding ASD symptomatology is the Salience Network (SN),^13^ which is believed to be integral in guiding attention to the most salient interoceptive and exteroceptive stimuli.^14^ Altered SN connectivity can discriminate children with ASD from neurotypical controls with high classification accuracy,^15^ and is associated with symptoms of restrictive/repetitive behaviors^15^, including atypical sensory processing^16^. Furthermore, recent work has found sustained hyperconnectivity with sensory processing regions within the SN throughout adolescence in youth with ASD^17^, which may highlight a potential neural mechanism underlying increased perceived salience of low-level, perceptual contingencies in the environment at the expense of higher-level social information^18^. The developmental origins of atypical SN connectivity in the context of ASD risk have not been previously examined, despite behavioral evidence this network in the emergence of ASD symptomatology in early infancy. Faces represent a highly salient class of stimuli for typically-developing infants^19^ and normative patterns of early SN connectivity may support the initial attentional bias toward faces as well as subsequent developmental increases in visual attention to faces in the first year^20^. Conversely, early disruptions in SN connectivity may iteratively derail processes that typically reinforce the perceived salience of faces^21^ by conferring heightened salience to lower-level non-social stimuli^18^. Indeed, initial overt symptom-based markers of ASD suggest deviations in processes that typically guide social communicative development, including attention to faces^5^ and speech. Potential atypicalities in SN connectivity may ultimately contribute to the emergence of characteristic features of ASD (i.e., social communicative impairments and altered sensitivity to sensory stimuli.

To test this model, we evaluated SN connectivity in 6-week-old infants at high (HR) and low (LR) familial risk for ASD and its association with subsequent behavioral ASD symptom-based markers, including atypicalities in visual social attention to faces, communicative development, and sensory processing. Between 4-8 weeks, early social-orienting behaviors transition from being under reflexive subcortical control to experience-dependent cortical control.^22^ Thus, examining SN connectivity during this period may reveal altered development of social attention and provide a mechanistic account for the ontogeny of ASD symptomatology. We hypothesized that HR infants would show SN hyperconnectivity with sensorimotor regions^16,23^ and that patterns of SN connectivity would predict individual trajectories of social attention, communicative development, and sensory sensitivities.

This study prospectively evaluated 53 HR and LR infants, with risk status based on family history:^24^ HR infants had at least one older sibling with ASD (N=24) whereas LR infants (N=29) had no family history of ASD or any other developmental disorders. Infants underwent restingstate functional magnetic resonance imaging (rs-fMRI) at 6 weeks of age during natural sleep to examine SN connectivity. Infants’ eye movements were tracked at 3-, 6-, 9-, and 12-months of age while they viewed video stimuli of naturalistic social interactions (i.e., excerpts from *Charlie Brown* and *Sesame Street*^25^) to capture developmental trajectories in visual social attention to faces. We examined core autism symptomatology with the Autism Observation Scale for Infants (AOSI) and nonverbal social-communication with the Early Social Communication Scale (ESCS) at 12 months (see Table 1), as well as sensory sensitivity with the Infant/Toddler Sensory Profile (ITSP) from 6- to 12-months.

**Table 1.** Participant Demographics.

|  |  | LR<br>N=29 |  | HR<br>N=24 |  | P |
| --- | --- | --- | --- | --- | --- | --- |
|  |  | N | % | N | % |  |
| Sex |  |  |  |  |  | 0.38 |
|  | Female | 11 | 37.93 | 17 | 58.62 |  |
|  | Male | 18 | 62.07 | 12 | 41.38 |  |
| Race |  |  |  |  |  | 0.88 |
|  | White | 20 | 68.97 | 17 | 58.62 |  |
|  | Non-white | 9 | 31.03 | 12 | 41.38 |  |
| Family Income |  |  |  |  |  | 0.23 |
|  | Not Answered | 1 | 3.45 | 0 | 0.00 |  |
|  | <50K | 3 | 10.34 | 6 | 25.00 |  |
|  | 50-75K | 4 | 13.79 | 4 | 16.67 |  |
|  | 75-100K | 5 | 17.24 | 5 | 20.83 |  |
|  | 100-125K | 4 | 13.79 | 2 | 8.33 |  |
|  | >125K | 12 | 41.38 | 7 | 29.17 |  |
|  |  | Mean | (SD) | Mean | (SD) | P |
| Birth Weight |  |  |  |  |  |  |
|  | Pounds | 7.65 | 1.67 | 7.41 | 2.43 | 0.67 |
| Age at Scan |  |  |  |  |  |  |
|  | Weeks | 6.63 | 1.37 | 6.65 | 1.09 | 0.95 |
| Relative Motion |  |  |  |  |  |  |
|  | mm | 0.23 | 0.82 | 0.10 | 0.07 | 0.44 |
| Behavioral Scores at 12 months |  |  |  |  |  |  |
|  | Mullen ELC | 110.22 | 11.83 | 106.67 | 17.24 | 0.39 |
|  | ESCS IJA | 0.71 | 0.45 | 0.96 | 0.48 | 0.06 |
|  | ESCS RJA | 0.31 | 0.23 | 0.28 | 0.24 | 0.56 |
|  | AOSI Total Score | 4.12 | 1.83 | 4.71 | 3.07 | 0.97 |
|  | Sensory Sensitivity | 19.52 | 4.39 | 20.7 | 7.55 | 0.54 |

Consistent with prior work, we used a right anterior insula (rAI) seed to characterize SN connectivity across the whole brain^16,26^ (Figure 1). We first examined between-group differences in SN connectivity by risk status (Figure 2A). Compared to LR infants, HR infants showed stronger connectivity between the hub of the SN (i.e., the rAI) and sensorimotor regions, including bilateral precentral gyrus and left postcentral gyrus, thalamus, hippocampus, caudate and putamen. In contrast, compared to LR infants, HR infants showed weaker connectivity between the rAI and prefrontal regions associated with higher-order processing and attentional control, including right inferior and middle frontal gyrus (IFG, MFG), and anterior cingulate. Importantly, we found an inverse relationship in SN connectivity with sensorimotor and prefrontal regions across all participants (*r*=-.*43, p* = 0.001, 95% CI: 0.18-0.63); infants exhibiting the strongest SN connectivity with sensorimotor regions also showed the weakest connectivity with higher-order prefrontal regions (Figure 2B). This direct tradeoff between neural resources allocated towards sensorimotor processing versus social attention could thus explain the co-emergence of both core sensory and social ASD symptoms.

**Figure 1.**
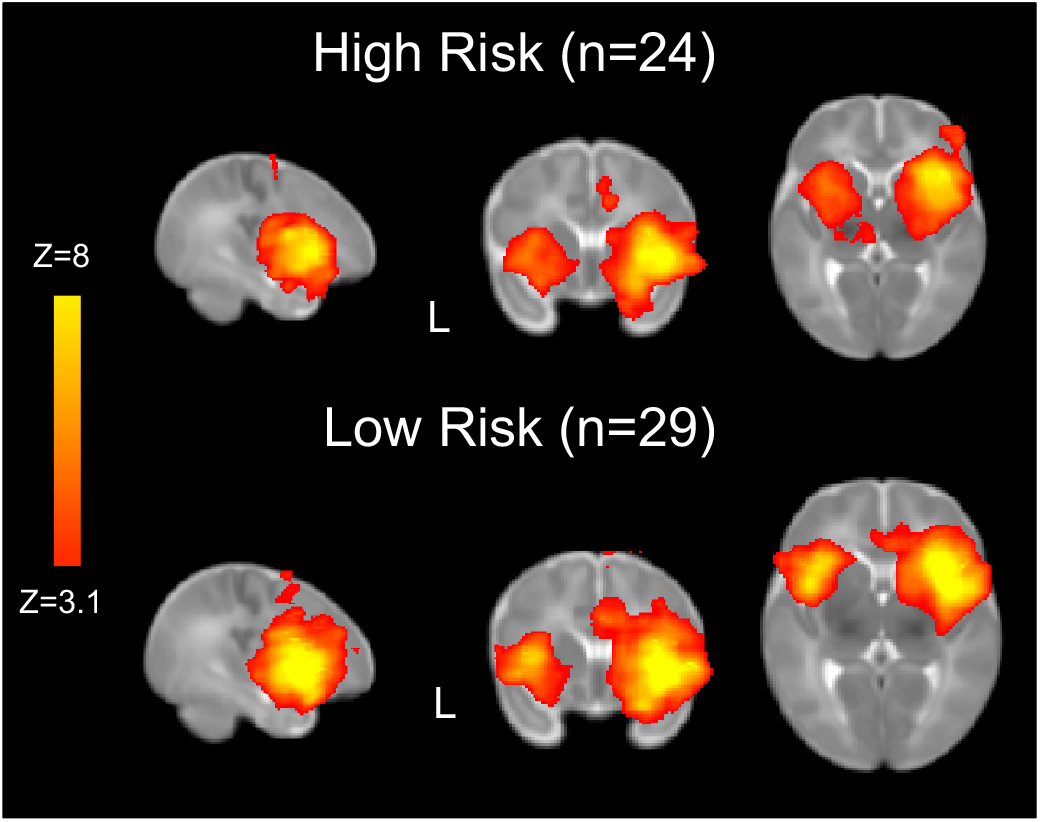
Salience Network maps. The Salience Network was detectable in both HR and LR infants using the right anterior insula (rAI) as the seed.

**Figure 2.**
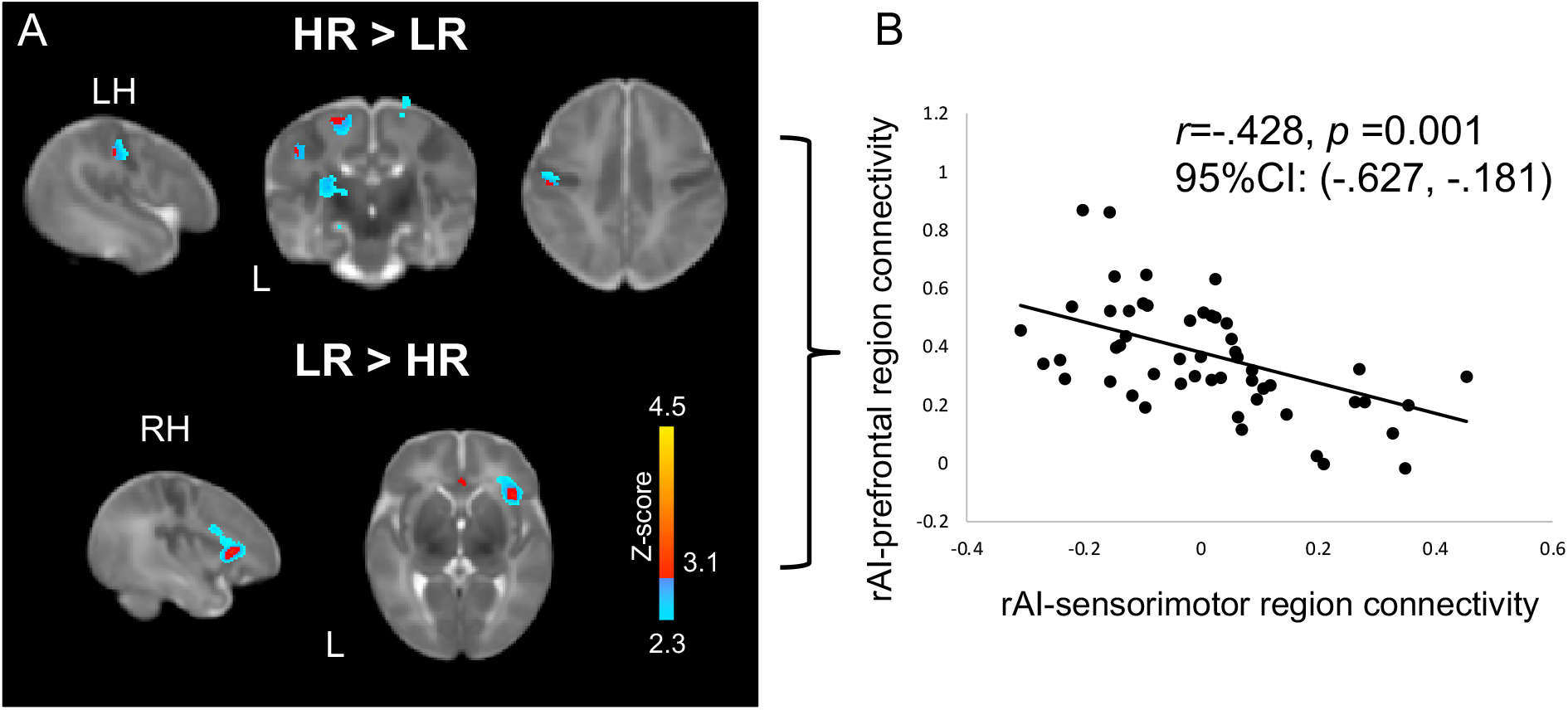
Between-group difference in Salience Network connectivity. *A*. Relative to LR infants, HR infants showed greater connectivity between the right anterior insula and left pre- and post-central gyri, thalamus, and caudate—regions associated with sensory and motor processing. In contrast, LR infants showed greater connectivity between the right anterior insula and right orbitalfrontal cortex and inferior frontal gyrus—frontal regions associated with social processing. B. Extracting connectivity z-scores from significant between-group clusters in SN connectivity revealed that these relative patterns of connectivity were inverse related with one another such that greater connectivity between the rAI and sensory regions was associated with lower connectivity between rAI and frontal, higher-order cognitive processing regions. For visualization purposes we display patterns of connectivity at both stringent (*Z*>3.1) and more liberal (*Z*=2.3) thresholds.

The inverse pattern of connectivity between the SN and social vs. sensorimotor regions indicates an early neural mechanism that may lead to divergent developmental outcomes for HR infants. To then directly examine the downstream, developmental effects of SN connectivity on visual social attention to faces, which has been shown to be attenuated in HR infants,^6^ we evaluated whether early SN connectivity predicts individual trajectories in attention to faces from 3- to 12-months. We measured percent looking time to faces from the eye-tracking data and used a Bayesian hierarchical linear model to estimate each infant’s rate of increased attention to faces from 3- to 12-months. This estimate was then used as a covariate of interest in our analysis of SN connectivity.

HR infants did not show any significant associations between 6-week SN connectivity and subsequent trajectories in attention to faces. In LR infants, however, greater connectivity between the two major hubs of the SN – rAI and the anterior cingulate cortex (ACC)^14^ – and right lateral OFC at 6 weeks predicted greater increases in attention to faces across the first postnatal year (Figure 3C). Importantly, these frontal regions predicting increased social attention overlapped with those exhibiting greater SN connectivity in LR infants relative to HR infants. Early social behavior, including orienting to faces and preferential attention to biological motion, is highly phylogenetically-conserved^3^ and genetically constrained^27^. Similarly, developmental changes in SN connectivity during the first postnatal year appears to be strongly influenced by genetic effects, more so than environmental factors^28^. In light of our findings, strong SN connectivity with frontal regions at 6 weeks may thus contribute to the early perceived salience of faces in typical development and thereafter support normative gains in social attention through iterative processes that further consolidate brain-behavior connections.

**Figure 3.**
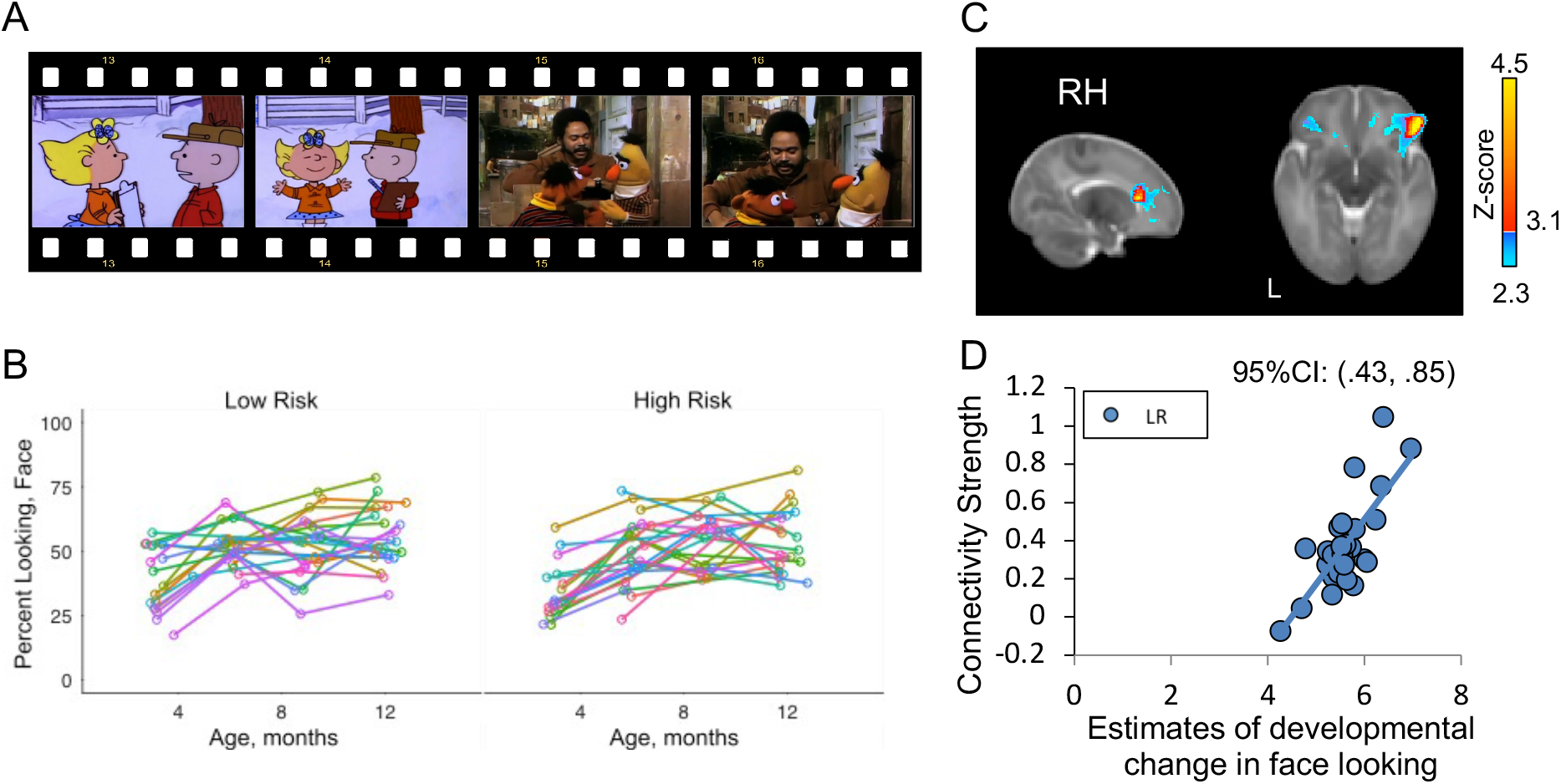
Salience Network connectivity and developmental trajectories in attention to faces. A. Example stills from the free-viewing eye-tracking stimuli. Each clip was approximately 2-minutes long and matched in color and motion. Each infant’s raw percent looking at faces across time points are plotted in B; individual data points were analyzed in a Bayesian hierarchical linear model. Estimates of each infant’s change in looking time to faces across age was derived from the best-fit model and included as a regressor of interest in a linear model of SN connectivity. Relative to HR infants, greater connectivity between the rAI and anterior cingulate cortex and right lateral orbitofrontal cortex was associated with greater increases in attention to faces from 3-to12-months of age. Estimates of connectivity strength in C and estimates of age on face-looking are plotted in D.

We next examined the relation between SN connectivity and standardized measures of communicative development (ESCS), ASD-related symptoms (AOSI), and sensory processing atypicalities (ITSP). In LR infants, greater connectivity between the SN hub (i.e., rAI) and both prefrontal (i.e., right IFG, MFG, and OFC) and subcortical regions associated with reward and learning (i.e., bilateral caudate, pallidum, and nucleus accumbens) predicted higher rates of initiating joint attention – nonverbal communicative behaviors associated with better language functioning and social competence^29^– at 12 months (Figure 4A). This finding provides further evidence that normative patterns of SN connectivity in early infancy support attention to socially relevant stimuli, thereby scaffolding the development of social communication skills.

**Figure 4.**
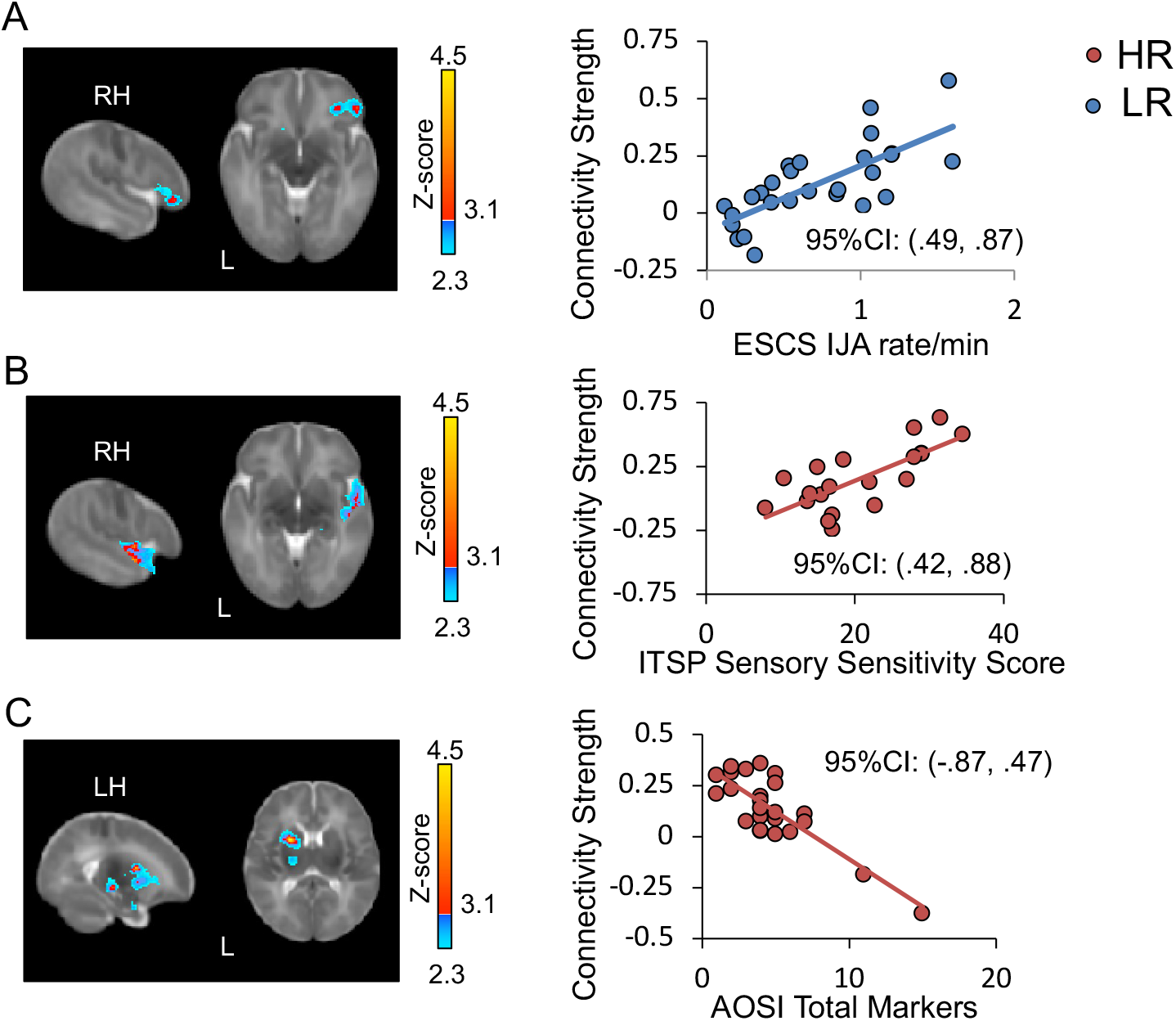
Associations between early Salience Network connectivity later social and sensory development. A. Greater connectivity with right anterior insula and right orbitalfrontal cortex, reward-processing regions, and prefrontal regions predicted greater rates of initiating joint attention in LR infants at 12months (ESCS IJA=Early Social Communication Skills, initiating joint attention. B. Greater connectivity with the right anterior insula and right superior temporal gyrus, amygdala and thalamus at 1.5 months predicted greater level of parent-reported sensory processing atypicalities in HR infants (ITSP=Infant Toddler Sensory Checklist). C. Greater connectivity with the right anterior insula and left basal ganglia, thalamus, and amygdala at 1.5 months predicted lower level of social impairment on the AOSI at 12 months in HR infants (AOSI: Autism Observation Scale for Infants).

Distinct patterns of 6-week SN connectivity also predicted behavioral markers of ASD risk in the HR infants at 6-12 months (Figure 4B and 4C). Greater SN connectivity with regions associated with primary auditory and sensory processing (right superior temporal gyrus and thalamus) predicted higher parental ratings of sensory hypersensitivity at 6-12 months. This finding is consistent with observed associations between SN hyperconnectivity and sensory over-responsivity in older children with ASD.^16^ HR infants may be predisposed to attend to extraneous sensory inputs quite early in development at the expense of socially relevant information. Furthermore, stronger connectivity between the SN hub and regions involved in implicit learning and reward processing (left putamen, IFG, caudate, amygdala), and memory (left hippocampus) predicted fewer ASD risk markers on the AOSI at 12 months. Better integration between the SN and both reward and emotional systems (e.g., regions associated with the social brain^30^) may be protective for social cognitive development in HR infants; conversely, reduced SN connectivity with these social brain regions in early infancy may negatively impact social learning.

Our data reveal dissociable patterns of SN connectivity between 6-week-old LR and HR infants with cascading effects on brain-behavioral processes that may underlie both normative and atypical social communicative development. Detectable differences in functional connectivity in early infancy corroborate cytoarchitectural and genetic data implicating prenatal neural development as a neurobiological risk factor for ASD.^31^ The observed hyperconnectivity between the hub of the SN and sensorimotor regions in HR infants likely represents a developmental vulnerability that, given the heterogeneity of developmental trajectories associated with familial risk for ASD, may be broadly associated with suboptimal outcomes. The observed tradeoff in functional connectivity with these areas versus higher-order prefrontal regions highlight that ASD risk manifests as differences of degree, not kind.

Taken together, these results provide clear empirical support for recent theoretical frameworks positing that initial deviations in attentional biases and/or sensorimotor processing may account for the emergence of ASD-related behaviors by altering the experience-dependent brain changes that typically guide social development.^32^ Furthermore, our findings demonstrate that atypical functional brain connectivity is present as early as 6 weeks of age, long before detectable differences in behavior are observed in infants and toddlers at high-risk for ASD. This difference is notable because it suggests a protracted prodromal period for known symptoms associated with ASD risk to emerge, including early social difficulties, attentional difficulties, language delays, and socioemotional problems. While specific patterns of SN connectivity within the HR and LR infants predicted distinct levels of social functioning and sensory processing, HR and LR infants exhibited comparable behavioral profiles at 12 months, consistent with prior behavioral studies.^10^ By affecting attentional biases, initial differences in brain function may become compounded over development through individual experiences, highlighting the role of experience in shaping brain-behavior relationships and the opportunity to intervene to prevent the crystallization of atypical social communicative development.

Examining early functional brain connectivity can inform the impact of genetic and environmental risk factors on subsequent development by highlighting the intermediary neural networks that ultimately underlie behavior. For instance, recent work has suggested that prenatal exposure to maternal inflammatory markers negatively influence early SN connectivity in infants without familial risk for ASD^33^, which suggests an intricate interplay between genetic and environmental factors in fine-tuning brain-behavior connections and contributing to the individual variability in early social orienting behaviors. While a primary aim of the current study was to elucidate the role of SN connectivity in conferring risk for ASD, infants with a family history of ASD are also more likely to exhibit atypical development, including subclinical ASD symptomatology (i.e., broader autism phenotype), speech/language delays, global developmental delay, and ADHD during school-age years.^34^ Thus, the clinical implications of our findings extend beyond risk of an ASD diagnosis, providing a testable model for examining how early SN connectivity may be related to other suboptimal developmental outcomes that are associated with familial risk for ASD. Although sample size and the single timepoint for evaluating SN connectivity are limitations in the current study, the overall results strongly suggests that atypical patterns of SN connectivity may reflect a developmental vulnerability. This is a possibility that should be examined in large-scale longitudinal studies that heavily sample brain and behavioral measures during the first postnatal years.

In sum, our findings demonstrate that aberrant patterns of functional brain connectivity can be detected in infants at high risk for developing ASD shortly after birth, and that these alterations predict ASD-related behaviors a year later. Identifying abnormal brain connectivity in infancy may pave the way for early interventions that can effectively redirect attention to social versus nonsocial inputs and thus promote development along normative trajectories.

## Methods

### Participants

Participants in this study were enrolled as part of a longitudinal project examining early brain-based markers of ASD during the first year. The Institutional Review Board (IRB) at the University of California, Los Angeles, approved all protocols associated with the project, and all enrolled participants had informed consent provided by their parent/legal guardian. All procedures complied with ethical regulations for vulnerable populations. Infants were assigned to risk-based cohorts based on family history: high-risk infants (HR) had at least one older sibling with a clinical ASD diagnosis whereas low-risk infants (LR) had no family history of ASD or any other developmental disorder. Prior research showed that the recurrence risk for developing ASD is approximately 20% in HR infants^35^. Exclusionary criteria for both groups included: 1) indication of genetic or neurological conditions associated with ASD risk (e.g., fragile X syndrome, epilepsy, tuberous sclerosis), 2) significant perinatal insult or chronic medical conditions impacting development, 3) severe visual, hearing, or motor impairment, 4) non-English speaking parents, and 5) contraindication for MRI (e.g., metal implants). All participants were enrolled in the study prior to 6 weeks of age. HR and LR infants were matched by gender (Mann-Whitney *U* = 306, *Z*=0.87 *p*=0.38, *r*=0.12), and birth weight (*t*(51)=0.43, *p*=0.67, Cohen’s *d*=0.12), as well as ethnicity and family socio-economic status (race: Mann-Whitney *U*=341.5*, p*=0.88; household income: Mann-Whitney *U*=282.5, *Z*=1.21 *p* = 0.23, *r=* 0.166). HR and LR infants did not differ on cognitive development at 12 months according to the Mullen Scales of Early Learning—Early Learning Composite (HR_ELC_ Mean=106.67, SD=17.24; LR_ELC_ Mean=110.22, SD=11.83; *t*(49)=0.87, *p* = 0.39, Cohen’s *d* =0.24; see Table 1).

A total of 53 infants (N=24 HR, N=29 LR) underwent fMRI during natural sleep at approximately 6 weeks of age (HR_age_ Mean=6.63 weeks, SD=1.37 weeks; LR_age_ Mean=6.65 weeks, SD=1.09 weeks, *t*(51)=0.062, *p*=0.95, Cohen’s *d*=0.02). A subset of 51 infants (N=24 HR, N=27 LR) provided data from behavioral measures of social and cognitive development at 12 months (2 LR infants dropped out of the study). Of the 53 infants, 50 infants (N=24 HR; 26 LR) provided longitudinal eye-tracking data at 3-, 6-, 9-, and 12-months of age. An additional 3 infants participated in the study but were not included in the analyses due to excessive head motion during scanning and/or scanner artifacts. No statistical methods were used to pre-determine sample size, but our sample size is larger than what has been reported in previous publications evaluating resting-state connectivity in early infancy ^36–38^

Whereas HR infants were all later-born children, the LR infants in our sample included both first-born (N=16) and later-born children (N=13). We did not observe differences in overall connectivity strength within the SN between first- and later-born LR infants (t(27)=1.07, *p*=.318, Cohen’s d = .379), nor did we observe differences in behavioral measures of cognitive and social development at 12 months (*t*’s<1.90, *p*’s>.06, Cohen’s *d* <0.53). Therefore, we do not believe birth order would have had an effect on the findings presented here.

### Behavioral Measures

Infants were administered a battery of behavioral assessments during the first postnatal year to measure socio-communicative, cognitive, and sensory development (see Table 1). At 12 months, infants’ developmental level was assessed on with the Mullen Scales of Early Learning^39^, nonverbal social communicative behaviors (i.e., rates of initiating of and responding to joint attention cues—IJA and RJA respectively) with the Early Social Communication Scales (ESCS^40^), and early signs of ASD symptomatology with the Autism Observation Scale for Infants (AOSI^41^). Approximately 20% of the total ESCS sample was double coded for reliability. Coders were trained undergraduate research assistants who were blind to ASD family risk status and other study variables. Intraclass correlations (ICC; absolute agreement, single measures) indicated good reliability for IJA (ICC = .96) and RJA (ICC = .89). Behavioral measures were administered blind to the infant’s risk status. Parents also completed the Infant/Toddler Sensory Profile,^42^ a standardized questionnaire tracking their child’s sensitivity to sensory inputs and sensory-related difficulties, at 6, 9, and 12 months; the average raw score on the Sensory Sensitivity quadrant was calculated and used as a general metric of sensitivity to visual, auditory, and tactile stimuli.

### Eye-tracking Protocol

Infants were eye-tracked at the 3-, 6-, 9- and 12-month visits while they were presented with two, 2-mintue full audiovisual video segments taken from a cartoon and live-action video; these video stimuli have been previously used in studies on visual social attention to faces in typically-developing infants.^25,43^ Infants sat on their caregiver’s lap during the eyetracking procedure at approximately 60 cm from the 65-cm video display monitor. Caregivers were explicitly instructed not to distract their infant’s attention from the screen during stimuli presentation.

Point-of-gaze (POG) data were collected using a Tobii T60XL eye-tracker at 60 Hz with a spatial accuracy of approximately 0.5° accuracy. Eye-movements (e.g., fixations, blinks, and saccades) were detected using the accompanying Tobii software. Infants’ POG were calibrated using a 5-point calibration scheme prior to data collection. The calibration scheme was repeated until an infant’s POG was within 1° of the center of the target and repeated between the two trials. The video stimuli were presented only after the calibration criterion had been reached. Individual trials were removed from analyses due to failure to initially calibrate the infant’s eyes to the eyetracking system (N_trials_=9) or failure to track an infant’s eyes because of excessive movement or fussiness (N_trials_ =31).

Video frames were 8-bit color images and 720 by 480 pixels in resolution. Each frame was hand-traced for areas of interest (AOIs), which were demarcated as a box encompassing each character’s face as in the prior studies of typical development using the same stimuli.^25,43^ Fixations that fell within AOIs were identified using software written in MATLAB (MathWorks, Inc; Natick, MA). The primary dependent measure was percent of fixations that were directed at the face AOIs.

### Eye-tracking Statistical Analysis

Our primary interest from modeling the eye-tracking data was to estimate each infant’s rate of change in attention to faces from 3 to 12 months, such that individual developmental trajectories in social attention could be analyzed as a function of 6-week SN connectivity. Longitudinal changes in percent fixation to faces were analyzed with a Bayesian hierarchical linear model in R (rstanarm package), which uses a Markov chain Monte Carlo simulation to draw a posterior distribution (e.g., a range of probable values for a variable given the data). This aspect of the Bayesian framework allows for greater precision in estimating parameters than by the frequentist approach.^44,45^

Attention to faces was operationalized as percent fixations to characters’ faces in each video stimulus. Developmental trajectories were modeled as the linear and quadratic effects of age (i.e., age and age^2^, respectively). Risk status (HR versus LR) and video stimulus type (*Charlie Brown* versus *Sesame Street*) were modeled as fixed effects; age and intercept were modeled as random effects to account for individual differences and correlated repeated measures at 3, 6, 9, and 12 months. The inclusion of the change of slope (i.e., the quadratic term age^2^) aimed to capture the change in growth rate from 3 to 12 months. Student-t distributions were used as priors for the regression coefficients and standard deviation, and a Gaussian function was used as the identity link function.

The model in equation form is:

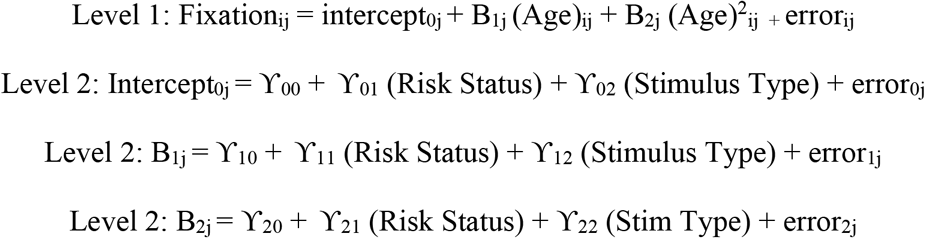

Pareto k diagnostic values indicated a good model fit (all pareto k estimates were less that 0.7). Diagnostics regarding model fit and visualization of posterior distributions were verified with the shinystan package in R. Statistical significance was evaluated using two-tailed tests. As expected, there was a significant linear effect of age such that attention to faces increased with age across all participants (95% CI: [1.73 9.39]). The rate of change overtime decreased across all infants indicating a quadratic trajectory in face-looking (95% CI: [−0.47 −0.01]). There was also a main effect of risk status; HR infants overall attended less to faces than LR infants (95% CI: [−40.91 −3.66]).

Using the coef function in R, we extracted each infant’s estimate of developmental increase in face-looking from 3 to 12 months (i.e., each infant’s beta coefficient for the age term in the model). These estimates were then used as a covariate of interest in a model of SN connectivity.

### MRI Data Acquisition

MRI data were acquired on a 3T Siemens Tim Trio scanner using a 12 channel head coil during natural sleep. Parents were instructed to put their infant to sleep using their normal bedtime routine. After the infant was asleep and swaddled, silicon earplugs were placed over the infant’s ear canal, and mini earmuffs were fitted over the entire outer ears. Infants were then placed on a custom-made bed that fit inside the scanner’s head coil and secured on the scanner bed with a Velcro strap. To minimize movement, a weighted blanket was used and foam pads were positioned around each infant’s head.

A localizer scan was used for graphic prescription. Structural matched bandwidth T2-weighted high-resolution echo planar images are acquired co-planar to the functional scans to ensure identical distortion characteristics to the fMRI scans (TR=5000ms, TE=34ms, matrix size 128×128, FOV=192mm, 34 slices, 1.5mm in-plane resolution, with 4mm-thick axial slices). Resting-state data were collected during an 8-min rs-fMRI scan (TR=2000ms, TE=28ms, matrix size 64×64, FOV=192mm, 34 slices, 3mm in-plane resolution, with 4mm-thick axial slices).

### fMRI Data Preprocessing

Functional imaging data were preprocessed and analyzed using FSL version 5.0.8 (fMRIB’s Software Library.^46^ Functional images were co-registered to the subject’s corresponding T2-weighted high-resolution anatomical scan, registered to an infant brain template^47^ using 12-parameter affine transformations, and spatially smoothed (Gaussian kernel of 6mm FWHM) to increase signal-to-noise ratio. ICA-AROMA was used to detect and remove motion artifacts from the data.^48^ This is a validated procedure that uses probabilistic independent component analysis to automatically detect participant-specific motion-related independent components while preserving signal of interest.^49^ ICA-AROMA was selected over other motion denoising methods, such as deleting individual motion-contaminated volume (e.g., scrubbing)^50^ to effectively control for motion while maximizing data from the full scan. HR and LR infants did not differ in the average number of noise components identified by ICA-AROMA [HR: Mean=27.25, SD=9.43; LR: Mean=27.72, SD=11.37; *t(51)*=0.16, *p*=0.87, Cohen’s *d* =0.05]; the number of noise components detected were comparable to that reported by Pruim and colleagues (23.1 components).^48^ HR and LR infants also did not differ on average relative motion prior to the denoising with ICA-AROMA (HR: Mean=0.10mm, SD=0.07mm; LR: Mean=0.23 mm, SD=0.82mm; *t(51)*=0.81, *p*=0.42, Cohen’s *d* =0.22). Data were then band-pass filtered (0.01Hz—0.1Hz). Nuissance regressors (e.g., mean cerebrospinal fluid, white matter, and global time series) from the bandpass filtered data were calculate and then regressed out of the filtered data to further remove noise. Given continuous debate in the field as per the pros and cons of implementing global signal regression^51^, group-level analyses were also completed without regressing global signal or its derivatives as nuisance variables; our were not were not affected by this change, including the inverse pattern of connectivity between frontal and sensorimotor regions (*r*=-.428, *p*=.001, 95% CI: [−0.626 −0.178]).

### fMRI Statistical Analyses

Resting-state fMRI analyses were conducted with FSL fMRI Expert Analysis Tool (FEAT, version 6.0 www.fmrib.ox.ac.uk/fsl/^52^). Whole brain connectivity within the SN was examined using a right anterior insula (rAI) seed^26^ that was derived from an anatomical parcellation of the right insula from a neonatal template. We identified the center of gravity of the right insula and created an anatomical mask composed of voxels anterior to the midline.^47^ The center of gravity for our rAI seed (X=23, Y=5, Z=1) corresponds to meta-analytic definitions of rAI^53^. Region-of-interest (ROI) time-series from each infant’s processed residuals in standard space were extracted and correlated with every other voxel in the brain to generate SN functional connectivity maps. Individual SN maps were converted into z-statistic maps using Fischer’s r-to-z transformation. At the group level, we modeled a 2-sample mixed-effects design, at *Z*>3.1 with cluster correction for multiple comparisons at p<.05, using FSL FLAME (FMRIB’s Local Analysis of Mixed Effects State) Stage 1+2. Statistical significance was evaluated using two-tailed tests.. For the between-group comparisons and regression analyses (see below), significance was assessed voxel-wise at p<.05, controlling for multiple comparisons using cluster-level correction estimated by AFNI’s 3dClustSim (version 16.3.05) with 10,000 iterations at an initial cluster forming threshold of p<0.001 (*Z*=3.1), a mixed-model spatial autocorrelation function (ACF), and a joint [HR+LR] SN connectivity map. Family income and gestational age were examined as potential covariates; as none contributed significantly they were excluded from the final analyses. Regression analyses with estimates of increases in face-looking from 3 to 12 months, AOSI, ESCS, and Infant/Toddler Sensory Profile were restricted to (i.e., masked by) the joint [HR + LR] SN connectivity maps (see Table 2). Parameter estimates of connectivity strength for between-group comparisons were extracted from significant clusters using FMRIB and evaluated using Pearson correlation.

**Table 2.** Coordinates of regions with significant functional connections to the right anterior insula.

| Region | Side | Peak mm (x,y,z) | Max Z |
| --- | --- | --- | --- |
| SN connectivity map between group comparison: HR>LR |  |  |  |
| Precentral Gyrus | L | (-14, -17, 40) | 3.41 |
| Postcentral Gyrus | L | (-35, -15, 27) | 3.25 |
| SN connectivity map between group comparison: LR>HR |  |  |  |
| Inferior Frontal Gyrus | R | (26, 11, 4) | 3.58 |
| Positive correlations between right anterior insula and Face-looking in LR group |  |  |  |
| Orbitofrontal Cortex | R | (35, 21, -5) | 4.61 |
| Inferior Frontal Gyrus | R | (29, 8, 7) | 4.36 |
| Anterior Cingulate Gyrus | R | (11, 13, 10) | 4.08 |
| Positive correlations between right anterior insula and ESCS IJA in LR group |  |  |  |
| Orbitofrontal Cortex | R | (33, 19, -4) | 3.25 |
| Positive correlations between right anterior insula and ITSP Sensory Sensitivity in HR group |  |  |  |
| Superior Temporal Gyrus | R | (35, -4, 8) | 3.85 |
| Negative correlations between right anterior insula and AOSI Total Markers in HR group |  |  |  |
| Putamen | L | (-14, 0 11) | 4.10 |
| Insula | L | (-19, 0, 10) | 3.41 |
| Hippocampus | L | (-16, -16, -2) | 3.41 |

### Additional Control Network Analyses (see Supplemental Materials and Supplemental Fig.1)

To address the specificity of our findings to the SN, additional analyses were conducted using posterior cingulate cortex. bilateral dlPFC, and bilateral pre/postcentral gyri seeds to separately evaluate connectivity in the default mode network (DMN), the fronto-parietal attentional network, and sensorimotor networks, respectively. Similar to our SN analyses, these seeds were derived from an anatomical parcellation from a neonatal template and identical analytic procedures were followed. We did not observe between group differences in patterns of connectivity within the DMN, fronto-parietal attentional network, or sensorimotor networks, which indicates specificity of our observed results for disruptions in SN connectivity in infants with a family history of ASD. In light of the null between-group differences, we did not run bottom-up correlations with behavioral measures of social functioning and sensory responsivity.

### Code Availability

Analysis of resting-state data were performed with software written in bash, and eye movements and coding of fixation data were conducted with software written in MATLAB (MathWorks), available upon request from the corresponding author.

### Data Availability

The data that support the findings of this paper are available via the National Database for Autism Research (NDAR; https://www.re3data.org/repository/r3d100010717).

## Acknowledgements

This work was supported by grants from National Institute of Child Health and Human Development (NICHD P50 HD055784, F31HD090937). The content is solely the responsibility of the authors and does not necessarily represent the official views of the National Institutes of Health. The authors are grateful for the generous support from the Brain Mapping Medical Research Organization, Brain Mapping Support Foundation, Pierson-Lovelace Foundation, Ahmanson Foundation, Capital Group Companies Charitable Foundation, William M. and Linda R. Dietel Philanthropic Fund, and Northstar Fund. The authors also wish to thank the infants and their families for their time and participation in the study, Carolyn Ponting and Rosemary McCarron for their assistance in project coordination and data collection, and the Child and Adult Neurodevelopmental Clinic for their contribution in behavioral characterization of the sample.

## Author Contributions

T.T. and M.D. developed the initial aims and design of the study. J.L. and T.T. contributed to the data acquisition and statistical analyses. K.L assisted in the processing and analyses of the data. T.T., S.G., S.J., S.Y.B., and M.D. interpreted the findings and contributed to the manuscript. T.T. wrote the initial draft of the manuscript and all authors revised and commented on drafts of the manuscript.

## Author Information

Authors do not have any competing financial interests to disclose. Correspondence and requests for materials should be addressed to

