## Supplemental Materials for "Altered Salience Network Connectivity in 6-Week-Old Infants at Risk for Autism"

Negative Control Network Analyses

Several seed-based functional connectivity analyses were conducted to evaluate the specificity of the observed between-group differences in Salience Network connectivity. The additional networks examined included the default mode network (DMN), the fronto-parietal network, and the sensorimotor network. Atypicalities in functional connectivity in the DMN and fronto-parietal networks have been observed in youth and adults with ASD, thus serving as good control networks to examine potential developmental origins of disrupted network connectivity in individuals with ASD [1–3]. However, the DMN and fronto-parietal networks undergo substantive development during the first year whereas the somatosentory network is present at birth [4]. Accordingly, examining this network allows us to further evaluate whether the observed between-group differences in network connectivity between HR and LR infants are specific to the SN or are more broadly reflective of disruptions in early functional brain organization[5].

These additional seed-based connectivity analyses were implemented according to the procedures described in the main text Methods section. Anatomically-defined seeds were used to identify the DMN, fronto-parietal network, and sensorimotor network. This included the precuneus (MNI: [-1, 30, 12]; Fox & Raichle, 2007) bilateral dlPFC (MNI: [+/-30,-10, 24]), and bilateral pre- and post-central gyri seeds (MNI: [+/- 11, 18, 1]), respectively. FSLMATHS [7] was used to identify the center of gravity (i.e., coordinates in MNI space) from an infant anatomical atlas [8] and create 5 mm seeds for each ROI for whole-brain seed-based connectivity analyses. The precuneus and pre-/post-central gyri are included as independent regions in the infant anatomical atlas we used; their parcellation did not include a separate dlPFC region of interest.

We created the dlPFC seed using the meta-analytic database on Neurosynth (https://neurosynth.org) [9] to identify MNI coordinates that correspond to the ROI.

As shown in Supplemental Figure 1, we did not observed any significant between-group differences in connectivity across the three control networks, attesting to the specificity of our findings of early atypicalities in Salience Network connectivity in infants at high familial risk for ASD.

*Equivalence Tests*

To further quantify the non-significant between-group differences in connectivity within the control networks, we conducted equivalence tests using the TOSTER package in R [10]. First, we used the FSL command fslmeants to separately extract parameter estimates of the whole-brain network connectivity for HR and LR groups, masked by the respective within-group connectivity map for each of the control networks (DMN, frontoparietal, and sensorimotor networks). As effect sizes are not commonly reported in resting state literature, we calculated the standardized effect size associated with a Z score >=3.1 and our sample size, which can be interpreted as the critical effect size. We used this value as the smallest effect size of interest (SESOI). Using this value, an equivalence test can reject effect sizes falling outside that bound. In this case, the critical effect size was $d$=0.855.

The TOST (two one-sided test) procedure for Student's equivalence test for independent samples, with equivalence bounds of $\Delta L = -0.855$ and $\Delta U = 0.855$, revealed that the effect observed is statistically equivalent for both the frontoparietal network ($t(51) = -1.861$, $p = 0.0342$) and sensorimotor network ($t(51) = -2.183$, $p = 0.0168$), but not the DMN network ($t(51) = -1.286$, $p = 0.102$). Whereas the connectivity strength for the somatosensory and frontoparietal networks between HR and LR groups are statistically equivalent, the confidence interval surrounding the between-group difference in connectivity strength for the DMN extends beyond the equivalence bounds.

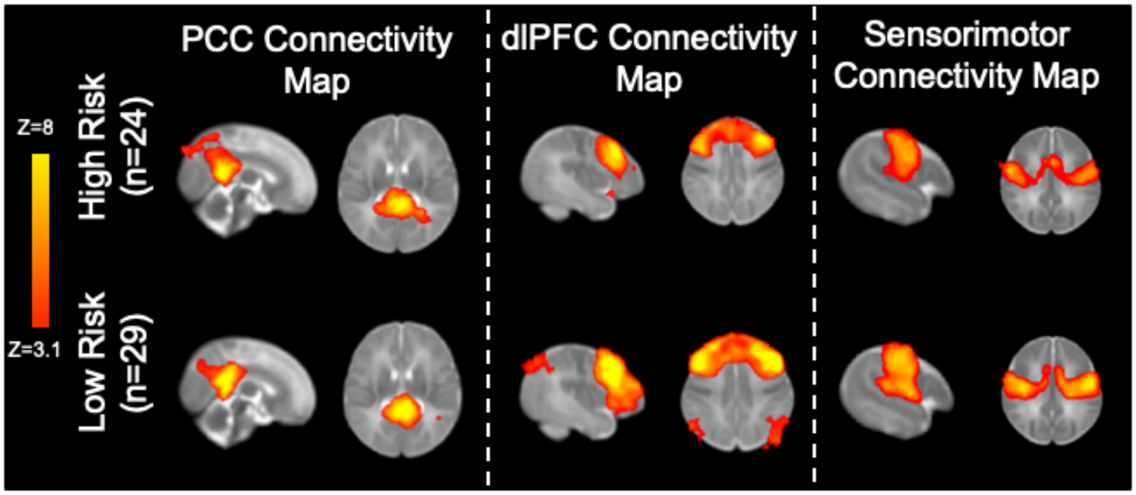


*Supplemental Figure 1.* Whole-brain connectivity maps using seed-based analyses from anatomically-defined PCC, dlPFC, and Sensorimotor seeds, respectively. We did not observe between-group differences
